# A MinION-based pipeline for fast and cost-effective DNA barcoding

**DOI:** 10.1101/253625

**Authors:** Amrita Srivathsan, Bilgenur Baloğlu, Wendy Wang, Wei Xin Tan, Denis Bertrand, Amanda Hui Qi Ng, Esther Jia Hui Boey, Jayce Jia Yu Koh, Niranjan Nagarajan, Rudolf Meier

## Abstract

DNA barcodes are useful for species discovery and species identification, but obtaining barcodes currently requires a well-equipped molecular laboratory, is time-consuming, and/or expensive. We here address these issues by developing a barcoding pipeline for Oxford Nanopore MinION™ and demonstrate that one flowcell can generate barcodes for ∼500 specimens despite high base-call error rates of MinION™. The pipeline overcomes the errors by first summarizing all reads for the same tagged amplicon as a consensus barcode. These barcodes are overall mismatch-free but retain indel errors that are concentrated in homopolymeric regions. We thus complement the barcode caller with an optional error correction pipeline that uses conserved amino-acid motifs from publicly available barcodes to correct the indel errors. The effectiveness of this pipeline is documented by analysing reads from three MinION™ runs that represent three different stages of MinION™ development. They generated data for (1) 511 specimens of a mixed Diptera sample, (2) 575 specimens of ants, and (3) 50 specimens of Chironomidae. The run based on the latest chemistry yielded MinION barcodes for 490 specimens which were assessed against reference Sanger barcodes (N=471). Overall, the MinION barcodes have an accuracy of 99.3%-100% and the number of ambiguities ranges from <0.01-1.5% depending on which correction pipeline is used. We demonstrate that it requires only 2 hours of sequencing to gather all information that is needed for obtaining reliable barcodes for most specimens (>90%). We estimate that up to 1000 barcodes can be generated in one flowcell and that the cost of a MinION barcode can be <USD 2.

## INTRODUCTION

DNA barcodes are widely used for species identification and discovery, but the existing methods for generating barcodes require well equipped molecular laboratories and there is a tradeoff between the cost of barcodes and the time needed for obtaining sequences. DNA barcodes obtained with Sanger sequencing are expensive, but can be obtained fairly quickly (Ivanova *et al.*, 2009) while barcodes obtained with Next Generation Sequencing (NGS) are cost-effective but require long processing time (Meier *et al.*, 2016; Shokralla *et al.*, 2014, 2015b; Hebert *et al.*, 2017; Wang *et al.,* 2018). In order to faciliate more widespread use of DNA barcodes for species discovery and identification, it is desirable to have techniques that require minimal equipment and yet generate DNA barcodes quickly and at low cost. Such techniques would democratize access to molecular identification techniques and make the methods more suitable for the numerous tasks that require fast identification of biological tissues (e.g., for pests, pathogens, vectors, illegally traded species, food ingredients: Ander *et al.*, 2013; Ball & Armstrong, 2006; Goncalves *et al.*, 2015; Shokralla *et al.*, 2015a; Tsui *et al.*, 2011).

Currently, most barcodes are still obtained with Sanger sequencing which requires access to a well-equipped molecular laboratory and an ABI sequencer. The literature is inconsistent about the cost of Sanger barcodes (Meier, 2008), but high throughput facilities like the Canadian Centre for DNA Barcoding charge C$2,200.00 per 96-well microplate (http://ccdb.ca/pricing/) which translates to ca. USD 17/specimen. Barcoding protocols based on NGS technologies such as Illumina (Meier *et al.*, 2016; Shokralla *et al.*, 2015b), PacBio SMRT (Hebert *et al.*, 2017) and Roche 454 (Shokralla *et al.*, 2014) have been described but they require expensive equipment, the sequencing run times are long, and such NGS barcodes are only cost-effective when large numbers of specimens are barcoded simultaneously.

We here test whether Oxford Nanopore Technologies (ONT)’s MinION™ is capable of delivering reliable and cost-effective DNA barcodes quickly without the need to have access to a well-equipped molecular laboratory. Introduced in 2014, the Oxford Nanopore Technologies (ONT) MinION™ sequencer is small and inexpensive (USD 900) and can be connected to a computer via a USB3.0-interface. The library preparation protocols are fairly simple and fast (10 min - 1.5 h) and the MinION™ allows for real-time data generation. This is particularly appealing when rapid identification is required for a biological sample (Borsting & Morling, 2015; Greninger *et al.*, 2015; Hoenen *et al.*, 2016; Kalianski *et al.*, 2015; Quick *et al.*, 2015; Quick *et al.*, 2016). However, what is much less desirable is MinION™’s low per base accuracy (∼90%: Hargreaves & Mulley, 2015; Ip *et al.*, 2015; Mikheyev & Tin, 2014; Sović *et al.*, 2016) which generates many bioinformatics challenges. These challenges are increasingly overcome through the development of new bioinformatic pipelines. For example, Loman *et al.* (2015) were able to reconstruct the genome of *Escherichia coli* with 99.5% nucleotide identity and MinION™ has also been used successfully for the identification of bacteria, plants, the characterization of microbiomes (Benitez-Paez *et al.*, 2016; Li *et al.*, 2016; Parker *et al.,* 2017; Shin *et al.*, 2016), and DNA fingerprinting (Zaaijer *et al.*, 2016).

Recently, MinION™ flowcells have also been used for generating barcodes for small numbers of specimens (N=7), but the existing pipelines are not cost-effective because barcodes are obtained in separate sequencing runs or by using Oxford Nanopore PCR barcoding kit (which can multiplex upto 96 samples) (Menegon *et al.*, 2017, Pomerantz *et al.*, 2017). We here propose an approach that allows for multiplexing hundreds of samples. It is a modification of our “NGS barcoding” pipeline that we initially optimized for Illumina platforms (Meier *et al.*, 2016; Wang *et al.*, 2018). In the original pipeline, we used tagged primers and a dual indexing strategy where 100 pairs of tagged forward and reverse primers can yield unique tag combinations for 10,000 specimens. Once such dual indexed products are generated, subsequent steps involve pooling of amplicons, DNA purification, preparation of DNA libraries, and sequencing.

The main challenge for a barcoding pipeline based on MinION™ is data analysis. This is the main reason why published studies on species identification have largely relied on mapping reads to known reference sequences (Benitez-Paez *et al.*, 2016; Shin *et al.*, 2016). However, this is often undesirable because the vast majority of eukaryote species lack barcodes (Meier *et al.*, 2016) which means that it often not known prior to a study whether a biological sample contains signals from species that have already been barcoded. We therefore here developed a pipeline that generates reliable barcodes without the use of of external references (i.e., use of reference barcode data is only optional). This pipeline was developed after testing existing tools. A new pipeline proved desirable after the test of existing tools revealed that they were not suitable for MinION™ data. For example, we initially attempted to use OBITOOLs (Boyer *et al.* 2015) for demultiplexing data but very few MinION™ reads could be demultiplexed due to high error rates in the tags. This problem can be overcome by using separate flowcells or ONT’s PCR barcoding kits (e.g., see Menegon *et al.* 2017, Pomerantz *et al.* 2017), but this makes MinION barcodes very expensive. In these studies the raw reads for one flow cell or library are assembled separately (e.g., using Celera (Myers *et al.*, 2000), or Canu (Koren *et al.*, 2017)) with error correction (e.g., Nanocorrect, Nanopolish (Loman *et al.*, 2015)). Note also that even when few samples were barcoded, the initial barcode estimates contained errors which Menegon *et al.* (2017) corrected by aligning reads to BLAST matches. Here we propose a pipeline that allows for generating up to 500 barcodes in one flowcell. We test this pipeline by analyzing data that were obtained in three flowcells using different taxa (ants, flies), scales (50->500 specimens/run), and MinION™ chemistry. We assess the reliability of MinION barcodes by comparing them to sequences obtained for the same PCR products with Sanger sequencing and Illumina MiSeq sequencing.

## METHODS

### PCR and Sequencing

We tested DNA barcoding using the MinION™ on three different sets of samples (Table 1). Our latest and main sample (A), comprises of PCR products from two pools of specimens: 254 specimens of dolichopodid flies (Diptera: Dolichopodidae) and 257 specimens from an assortment of flies (Diptera) from different families (hereon referred to as mixed Diptera sample). (B) Our second sample consists of PCR products for 575 ant specimens (Hymenoptera: Formicidae). (C) The final dataset consists of tagged amplicons for 50 chironomid midges (Diptera: Chironomidae). Note that the three datasets were obtained using different generations of flowcell and library preparation methods with A being the most updated dataset using the 1D^2^ gDNA sequencing kit (SQK-LSK308) running on a R9.5 MIN107 flowcell. For sample sets B and C, we obtained 2D reads.

**Table 1:**
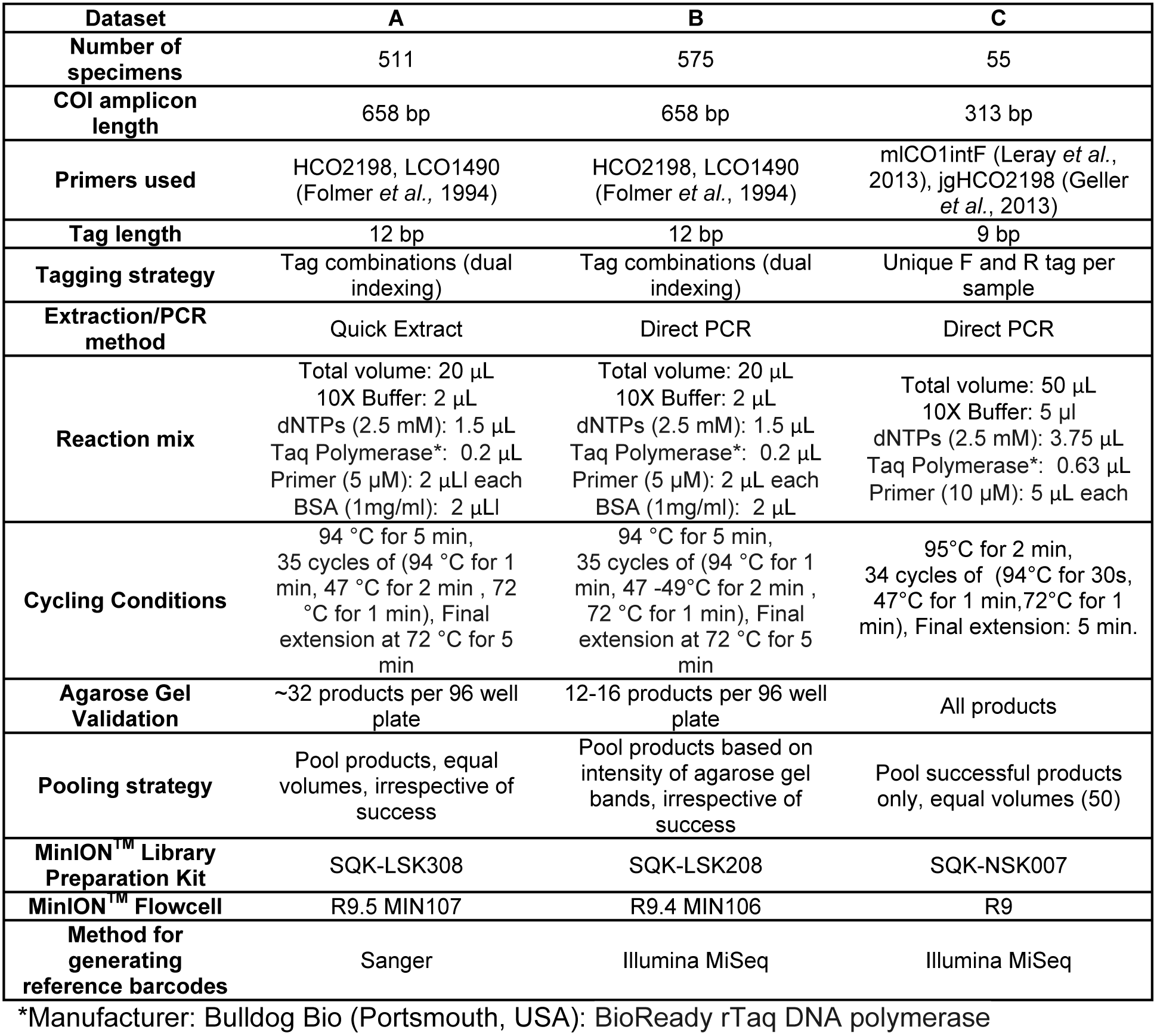
Experimental design for obtaining the three datasets

DNA barcodes were amplified either by ‘Direct PCR’ (for ants and chironomid midges) (Wong *et al.,* 2014) or using DNA extracted from whole fly specimens with 10 μL of QuickExtract (Epicentre, Wisconsin, USA). For sets A and B, PCR was used to amplify the 658 bp fragment of Cytochrome C oxidase subunit I (COI) (Folmer *et al.*, 1994). Each primer was tagged with a unique 12 bp sequence for the sequence to specimen association (Meier *et al.*, 2016; Wang *et al.*, 2018). The tags were designed using Barcode Generator (http://comaiwiki.genomecenter.ucdavis.edu/index.php/Barcodes) where minimum number of mismatches between tags was set at 6 bp. Moreover, it was ensured that no substrings of ≥6bp or more was shared between any two tags. Twenty five such tags were created, allowing for tagging 625 products using a dual indexing approach. For dataset C, we amplified a shorter 313 bp fragment using degenerate metazoan primers (Table 1). We used existing primers tagged with 9 bp sequences that are being routinely used for DNA barcoding using Illumina MiSeq/HiSeq (Meier *et al.*, 2016; Wang *et al.*, 2018). Given that these were short tags, it was ensured that both forward and reverse tags were unique for each specimen.

Pooled products (see Table 1) were first purified with SureClean™ (Bioline, London, UK). In some cases, an additional bead-based clean-up was employed to remove any remaining primer-dimer sequences. For this clean-up, we used 0.2% Sera-Mag Carboxylate-Modified Beads (GE Healthcare Life Sciences, Malborough, USA) in 18% PEG-8000 (polyethylene-glycol) solution at optimized DNA to Beads+PEG solution ratio (Rohland & Reich, 2012; Faircloth & Glenn, 2014). Purified products were used for library preparation using the MinION™ sequencer.

DNA concentration of the amplicon pool was determined using Qubit fluorometer 2.0 (Thermo Fisher Scientific, Waltham, USA). One microgram of amplified product was used for MinION™ library preparation using the SQK-LSK308, SQK-LSK208 and SQK-NSK007 library preparation kit for sample sets A, B and C respectively. Library preparation was carried out according to the manufacturer’s protocol with the omission of DNA fragmentation and DNA repair as amplicons were used as DNA input. Briefly, amplified product was end-repaired using NEBNext Ultra II End-Repair/ dA-tailing Module (New England Biolabs, Ipswich, USA) at 20°C for 5 min and 65°C for 5 min and then cleaned up with 1X AMPure XP beads (Beckman Coulter, Brea, USA). Adapter ligation used NEB Blunt / TA Ligase Master Mix (New England Biolabs, Ipswich, USA) together with the adapter provided in the kits. Ligation was achieved at room temperature (10 min). For sample set A, 0.4X AMPure XP beads (Beckman Coulter, Brea, USA) were used for clean-up before a second adapter ligation step was carried out. The adapted library was purified with ABB buffer provided in the SQK-LSK308 kit (Oxford Nanopore Technologies, Oxford, UK). For sample sets B and C, HPT was added and the ligation reaction was further incubated at room temperature for 10 min. Adapted DNA was purified using washed MyOne C1 beads (Thermo Fisher Scientific, Waltham, USA). The final library was then loaded on a flow cell and sequenced using the respective workflows on MinKNOW. Total library preparation time was estimated to be 2h. The libraries for samples B and C were split into two loads and loaded 24h apart. Sample set A was loaded into the flowcell in one load. 1D^2^ reads were base-called using Albacore (version 1.2.4) for data set A, 2D reads were base-called using Albacore (version 1.1.0) for data set B, and Metrichor (RNN SQK007 1.99) software with 2D Basecalling for SQK-NSK007 was used for data set C. The fastq files were generated by Albacore for data set A and poretools (version 0.5.1-17, option --type 2D) for data set B and C (see supplementary table 1, and supplementary figure 1 for read length distribution, supplementary figure 2 for read GC-content distribution and supplementary figure 3 for 5-mer frequency distribution).

In order to assess the quality of DNA barcodes obtained using the MinION™ sequencer, we sequenced the same PCR products using either Illumina MiSeq (for sample sets B and C), or using Sanger sequencing (for sample set A). For Sanger sequencing, PCR products were purified using SureClean Plus (Bioline, London, UK), and cycle sequencing was carried out using BigDye™ (ThermoFisher Scientific, San Jose, USA) under manufacturer’s recommendation. Cycle sequenced products were precipitated using PureSeq (Aline BioSciences, Woborn, USA) and analysed in an ABI 3730xl 96 capillary sequencer. For Illumina sequencing, libraries were prepared using TruSeq Nano kit (Illumina, San Diego, USA) and sequences were generated using MiSeq (300 bp, paired end). Note that for sample set B, products of 658 bp were sequenced using MiSeq, thus we obtained sequences of 300 bp at 5’ and 3’ ends of the sequences.

### Bioinformatic Analysis

#### Obtaining Reference barcodes with Sanger Sequencing and MiSeq

ABI sequences were edited and assembled in Sequencher v4 (GeneCodes, Ann Arbor, USA). For Illumina MiSeq of the 313 bp COI fragment (dataset C), we followed the bioinformatics procedures developed in Meier *et al.* (2016) to demultiplex and call the COI barcodes. This involved demultiplexing using a custom script that looks for perfect tag matches and allows for up to two substitution errors in the primers (Meier et al., 2016, Wang et al. 2018). To obtain the barcodes, the demultiplexed data are merged into unique sequences while retaining the count information. The count of the most dominant sequence (i.e. most abundant sequence) is then compared with the count of the next most dominant sequence. The dominant sequence is then accepted as the valid barcode for the specimen if it had a minimum 10X coverage and it was at least five times in frequency as the next most dominant sequence (i.e. ratio of sequence counts of subdominant to dominant sequence <=0.2). This latter criterion ensures that contaminated PCR products are not used to obtain reference barcodes. For dataset B where 658 bp products were sequenced using Illumina MiSeq, a different strategy had to be applied because 300 bp PE sequencing does not allow for the recovery of a full-length DNA barcode because approximately ∼133 bp in the middle of each barcode are missing. In order to demultiplex the data using Meier *et al.* (2016) pipeline, we initially merged the two reads end-to-end after fixing the orientation of read 2. After demultiplexing, the sequences were split back again as this initial merged sequence cannot be directly compared with MinION barcodes which are full length (658 bp). As there was a drop in quality scores beyond 200 bp for this dataset, we furthermore trimmed each end to 200 bp only. Following this, the same pipeline as for dataset B was applied and two 200 bp fragments of the COI barcode were obtained. Any specimen with either of the two fragments failing the 10X coverage or ratio criterion was excluded.

#### Analysis of MinION™ data

##### Primer identification and demultiplexing

Tools developed for Illumina data, such as OBITools (Boyer *et al.* 2015) can be applied to MinION™ data, but they perform poorly; for example only 2.3-7.5% of the MinION™ reads can be demultiplexed even if a large of number of errors are allowed for primers (e=5). Hence we had to develop a new pipeline (Table 2) which starts with identifying the primer in a MinION™ sequence (implemented in miniBarcoder.py). Subsequently, the barcode and tag are retrieved from the 3’ and 5’ ends of the primer respectively. Primers were aligned to the reads generated from the MinION™ using *glsearch36* (Pearson, 1990), where the primer is used as the query and the raw reads are used as the database. This allows for an alignment that is global (end-to-end) for the query and local for the reference sequence. Multiple gap and e-value parameters were tested for dataset A and we chose those parameters that yielded the consensus barcodes of the highest quality (for details, see supplementary Figure 4a). The performance of these parameters were then tested by also applying them to the data from dataset B and C and assessing the quality of the barcodes that were obtained with these parameters.

**Table 2:**
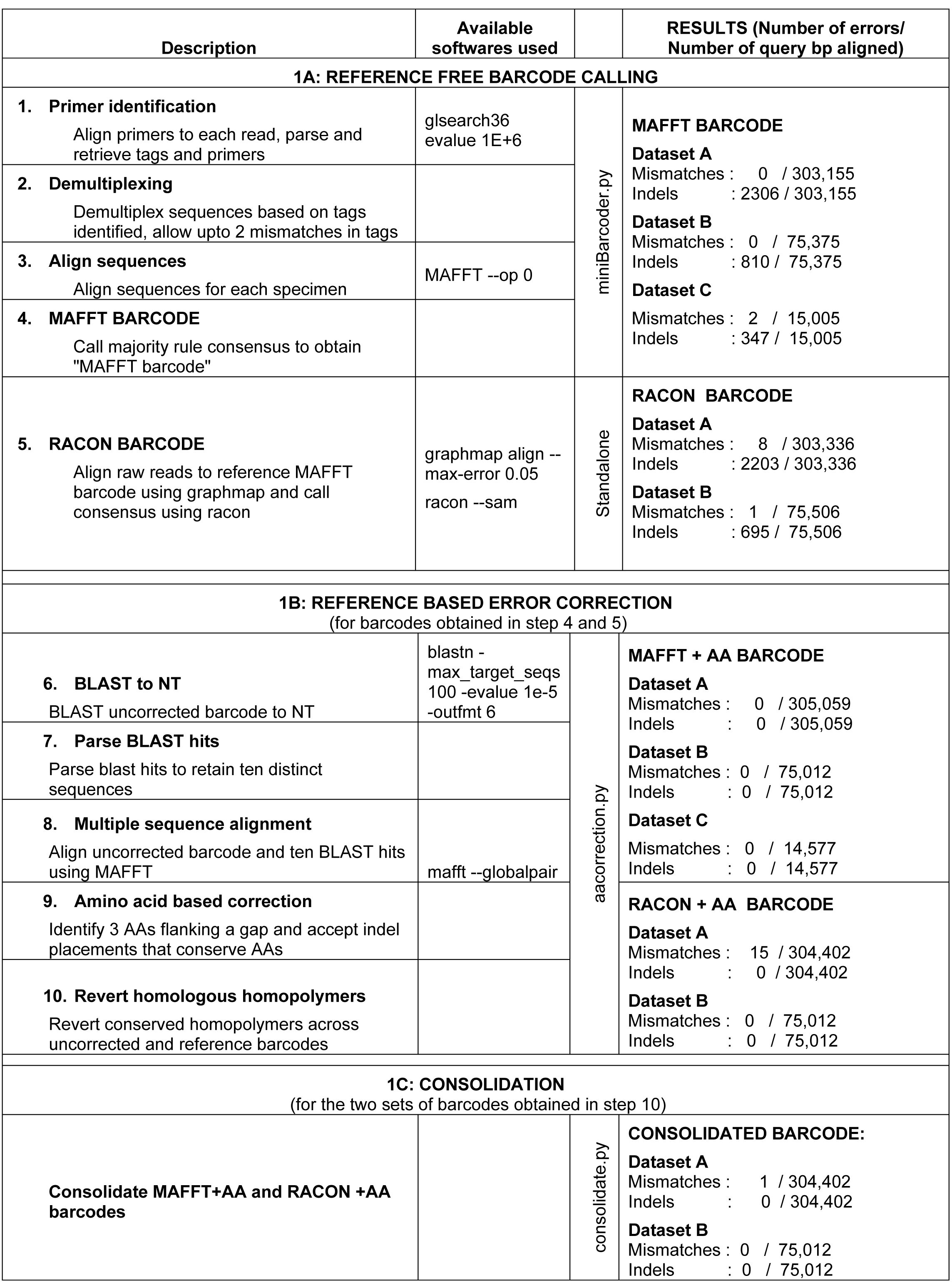
Key steps involved in the pipeline developed. Prior to using this pipeline base-calling should have been done and FASTA and FASTQ files generated.

Next, we retrieved the tag associated with each primer as the sequence flanking the 5’ end of the primer (hereon called “retrieved sequence tags”). The combination of forward and reverse tags could be used to demultiplex the data. The start and end site of the barcode was identified as the region between 3’ ends of the two primers. There are two reasons why we use two different processes for identifying primers and tags: 1) demultiplexing accuracy can be controlled by applying stricter parameters to the tags than primers; 2) the running time for analysis is shorter because there are only 2 primers (forward/reverse), but many tags (up to 25). In order to demultiplex reads, the sequence tags have to be matched to the original reference tag sequences that correspond to each specimen (see demultiplexing file). However, given the high error rates for MinION™ reads, the retrieved sequence tags often contain errors and need to be matched to the best-matching, original tag sequence. This was accomplished by “mutating” the original tag sequence so that all possible strings are generated that contain up to two errors (substitutions, insertions or deletions: see Supplementary Figure 4b for parameter choice). Note that only twenty-five 12 bp tags were used in our experiments so that it is not time-consuming to create these “mutant” strings. The set of original and imperfect reference tags are then matched against the retrieved sequence tags.

#### Obtaining MinION barcode hypotheses

Our pipeline starts by generating a first barcode hypothesis that is obtained via reference-free consensus sequence calling. Such barcodes can then be further refined with existing MinION™ tools (Racon) and/or by using a correction pipeline that takes advantage of information obtained from external barcode databases.

1. <u>MAFFT consensus barcodes</u>: The demultiplexing of the MinION™ data yields a read set for every amplicon. The reads in each set are aligned before a consensus barcode is called. For this purpose, identical reads are merged while retaining count information. Afterwards, the reads are aligned using MAFFT v7 (Katoh & Standley, 2013). Different alignment parameters were tested and we chose those that maximized congruence (mismatches/indels) between consensus barcodes based on MinION™ data and the sequences obtained with Sanger and/or Illumina sequencing; i.e., MAFFT was run under default (or auto) mode with gap-opening penalty set as 0 (multiple parameters were tested for MAFFT: --auto, --ensi, --globalpair,--genafpair; gap opening penalties via varying –op: default, 1, and 0). These settings generate a large number of gaps in the Multiple Sequence Alignment (MSA). Many of these indels are initially retained when the majority rule consensus barcode is called. However, we then delete these indels and score all positions lacking a base-call in >50% as ambiguous (“N”). This yields a first barcode hypothesis that we call “MAFFT barcode”. Comparison with Sanger and Illumina data revealed that MAFFT barcodes are mismatch-free, but retain indel errors. Note that one advantage of such an alignment-consensus approach is that contaminant PCR products can be readily identified because they have an unusually large number of ambiguous bases. For this study, we removed any sequence that had >1% of bases called as “N”s.
2. <u>Racon barcodes</u>. We also tested whether existing consensus calling tools for MinION™ data can yield better barcode hypotheses. We used one of the latest tools (“Racon”: Vaser *et al.* 2017, git commit 0834442) by first mapping all reads onto the barcode generated with MAFFT (Graphmap v0.5.2: Sović *et al.* 2016) and then processing the sam file with Racon (--sam). Initially this led to a number of substitution errors (>70 barcodes with mismatches in dataset A) but the performance improved considerably after removing the reads with the largest numbers of errors. This removal can be achieved through mapping reads onto the MAFFT barcode. We tested different identity settings for the read removal (--max-error) in Graphmap (default=disabled and 0.1 - 0.05). Eventually, we used Racon for analysing the 0.05 read set. This yielded barcodes with more mismatches but fewer indel errors compared to the original MAFFT barcodes.

Note that another way to obtain barcodes from binned raw reads would involve sequence assembly as implemented by Pomerantz *et al.* (2017) who use NanoFilt (https://github.com/wdecoster/nanofilt) and Canu (Koren *et al.* 2017). We tested this under the assemble mode with parameters suggested by Pomerantz *et al.* (2017) (minOverlapLength=50, genomeSize=1000, minReadLength=100), but this pipeline yielded few consensus barcodes and was then abandoned (see results).

#### Further Refinement of MinION barcode hypotheses through the use of external data

##### Indel errors

Consensus barcodes based on MinION™ reads contain indel errors because MinION™ makes “counting errors” in homopolymeric regions. For protein-encoding genes, many of these errors can be identified / corrected because they tend to cause frame shifts or changes in amino acid assignments. Both issues are best diagnosed by comparing MinION barcodes to barcodes from public databases given that COI sequences are protein-encoding and intron-free (see Supplementary Bioinformatics methods Figure 1). We therefore used BLAST to find 10 similar barcodes in NCBI (filtered to retain unique haplotypes). We then aligned the MinION barcode hypotheses (MAFFT and Racon) to these NCBI barcodes in order to identify the approximate location of indel errors in the MinION barcodes. Insertion errors were then corrected through deletion and gaps through replacement with ambiguous bases (“N”). However, standard alignment software only yields only one MSA even if the precise gap placement is ambiguous. In order to correct for the uncertainty in gap placement, our pipeline tests all placements within a three amino-acid window on either side of the indel placement in the initial MSA. We then identify those indel placements that yield conserved AA motifs when compared to the 10 best-matching NCBI barcodes. Note that due to strong stabilizing selection on the COI amino acid sequence (Kwong *et al.*, 2012), even poor-quality BLAST matches (80% identity) tend to yield conserved AA motifs. However, given that the genetic code is degenerate, there are usually multiple gap placements that conserve amino acid assignments. This means that usually multiple nucleotide sites have to be replaced with ambiguous bases and a single missing base in a MinION barcode hypothesis can lead to the insertion of multiple Ns.

This is undesirable, but inspection of the placement of ambiguous bases reveals that almost all are in homopolymeric regions. Many are 1^st^ and 2^nd^ positions that are conserved across all of the 10 best-matching barcodes obtained from NCBI (even if the identity is as low as 80%). Given that it is known that MinION™ reads introduce errors in homopolymer regions, we thus tested whether “Ns” in homopolymeric regions (>=2bp) can be replaced with homopolymeric bases from the BLAST hits as long as (1) the regions were conserved across all of the 10 best and unique hits and (2) the replacement was consistent with the MAFFT or Racon barcode. This replacement procedure also has an evolutionary justification given that conservation across all BLAST hits means that the sites have been conserved for millions of years (poorest BLAST hits for MinION barcodes often have <90% identity). It is thus not surprising that applying this correction did not introduce a single mismatch to the 471 barcodes for which we also have barcodes that were obtained with Sanger or Illumina sequencing.

##### Consolidation of MAFFT and Racon barcodes

MAFFT barcodes include next to no mismatch errors while Racon barcodes have fewer indel errors. It is thus advantageous to combine both after applying the AA correction pipeline. This fusion is accomplished by aligning the corrected MAFFT and Racon barcodes and calling the consensus. We find that there are few instances of mismatch between the corrected Racon and MAFFT barcodes and resolve these conflicts by accepting the MAFFT+AA solution given that MAFFT+AA barcodes contain no mismatch errors (see results).

##### Implementation

We created a pipeline (implemented in aacorrection.py, Table 2) that performs the following steps sequentially: First, the best BLAST hits for the MAFFT and Racon barcodes are found with MEGABLAST to NCBI’s “nt” database (e-value < 1e-5) and a FASTA file for sequences for the top hundred hits (locally or remotely) is retrieved. The hits are parsed to retrieve only the region of the sequences that overlaps with the query reference barcode in the correct orientation. Following this, ten distinct sequences that are most similar and most overlapping (five each) to reference barcode are retained and aligned (MAFFT, --globalpair). The correct reading frame is identified. Once an indel (< 5 bp) is encountered in the MinION barcode, a window corresponding to codon sequences of three flanking amino acids on either side of the indel is retrieved. In this window, depending on number of indel errors; all possible placements of the missing or additional nucleotides are assessed by checking whether it conserves the amino acid assignment. We then insert “Ns” into all placements that conserve AA assignments. Next we revert those N’s back to nucleotides that are in homopolymeric regions that are conserved across all BLAST hits. This correction is applied to both MAFFT and Racon barcodes. Afterwards, the AA-corrected MAFFT and Racon barcodes are consolidated (consolidate.py, Table 2). We allow the user to limit the procedure to sequences of certain length; in this study, we correct consensus sequences within 640-670 bp for full length barcodes and 300-330 bp for the mini-barcode.

Note that comparison with external data is also implemented in another recently developed pipeline (“ONtoBAR”: Menegon *et al.* 2017). It assembles the MinION™ reads in reference to a barcode from NCBI and then obtains a consensus. We tested this approach on a subset of barcodes (n=10, Supplementary Table 2) but found that it creates mismatch errors in low coverage regions (likely due to inclusion of reference bases). This was particularly common when distant NCBI barcodes were used.

### Validation

To validate the MinION barcodes, we compared each corrected MinION barcode with the corresponding reference barcode obtained with Illumina or Sanger. We aligned the MinION barcode with the reference barcodes using MAFFT (Katoh & Standley, 2013) and calculated the number of (a) mismatches, (b) gaps introduced and (c) ambiguous bases in the MinION barcode. For uncorrected data, indel errors can lead to overestimation of substitution errors using MAFFT and hence the errors were measured using dnadiff (MUMmer 3.0, Kurtz *et al.* 2004).

#### Assessment of sequencing biases in homopolymeric regions

In order to assess the sequencing errors in homopolymeric regions for MinION™, we calculated the number of pentamers of A/T and G/C for each demultiplexed barcode dataset. Frequency of pentamers was calculated as number of pentamers/total number of bases in the dataset. Similarly, we calculated frequency of pentamers for each MinION™ MAFFT consensus barcode and reference DNA barcode. We compared the distribution of frequency of pentamers in reference barcode and (a) MinION™ raw reads and (b) MinION™ consensus barcode. This analysis was limited to Dataset A, as this dataset has been generated using the latest MinION™ chemistry.

### Estimating species composition

All barcodes were aligned using MAFFT v7 (Katoh & Standley, 2013) under default parameters. We afterwards determined the number of species (or Molecular Operational Taxonomic Units (mOTUs)) using “objective clustering” as implemented in SpeciesIdentifier (Meier *et al.*, 2006) and by using a custom-built Python script (both using p-distances at various thresholds: Srivathsan & Meier, 2012). For these mOTU delimitations, we treated gaps as missing data.

### Effect of run time on sample characterization

We assessed the relationship between MinION™ sequencing run time and the number of reliable barcodes obtained. Read time stamps were used to generate datasets for each hour. Number of specimens demultiplexed, and coverage per specimen was determined for each hourly dataset. Moreover, MAFFT consensus barcodes and corresponding error-corrected consensus barcodes were obtained as described above. Given that we were handling up to 48 hours of data generation and ∼900 datasets, we did not apply Racon correction and consolidation. Following this, at each time-point, the number of mOTUs and their abundances were characterised in order to assess when the species composition in a sample can be determined reliably. Species composition at a given time point was compared with the final dataset using Bray-Curtis dissimilarity using *vegdist* in *vegan* package (Oksanen *et al.*, 2015) in R v3.2.1.

## RESULTS

The number of sequences obtained from the MinION™ sequencer varied across the three runs, with the latest run yielding 2,046,461 million 1D^2^ reads, while the older runs produced less data (Table 3). For sample A, we submitted 511 PCR products (254: Dolichopodidae and 257: mixed Diptera) and 17 negative controls for sequencing and managed to generate Sanger barcodes for 482 of the 511 PCR products (94.32% success). In order to demultiplex the MinION™ data, we tested different parameters in order to optimize the search for primer sequences and tag binning. The varied parameters included e-value, number of gaps allowed in the alignment of the primer, and number of mismatches allowed for the tags (Supplementary Figure 4). We eventually selected those parameters that allowed for high read recovery during demultiplexing while keeping the number of ambiguous base calls in the MAFFT consensus barcode low (Supplementary Figure 4). The chosen parameters were e-value=1e+6, maximum of 5 gaps for primer identification and 2 mismatches for tag identification. This yielded 294,887 demultiplexed reads thus ensuring at least 10X coverage for all 511 products (Table 3). None of the negatives had more than 10 reads thus suggesting accurate demultiplexing of data.

**Table 3:** From reads to species identification for the three datasets

| Datasets | A |  | B | C |
| --- | --- | --- | --- | --- |
| Taxon | Dolichopodidae | Mixed<br>Diptera | Formicidae | Chironomidae |
| MinION Output |  |  |  |  |
| Reads Total/Demultiplexed | 2,046,461/294,887 |  | 673,266/ 194,495 | 14,722/7,594 |
| Barcode Generation |  |  |  |  |
| Number of amplicons | 254 | 257 | 575 | 50 |
| Number of barcodes >10X<br>/<1%N's | 254/243 | 257/247 | 428/395 | 50/48 |
| Accepted barcodes | 243 | 247 | 358* | 47 |
| Species Richness Estimation |  |  |  |  |
| Number of 3% mOTUs<br>(MAFFT raw/corrected<br>barcodes) | 33/33 | 73/73 | 20/20 | 3/3 |

\* 1/359 sequences is bacterial

Applying the MAFFT consensus procedures to the 1D^2^ data yielded 490 barcodes (<1% of the bases called as Ns) with 243 corresponding to Dolichopodidae and 247 to mixed Diptera. The Canu assembly approach on the other hand yielded only 174 barcodes and was then abandoned. Comparison of MAFFT consensus barcodes with corresponding Sanger sequences (n=471), revealed the MAFFT barcodes had no substitution error (this excludes one sequence with a substitution error of >100 bp which is likely due to wet lab error). However all sequences contained an average of ∼5 indel errors (0.76%) per DNA barcode (Table 2,4). Polishing the MAFFT consensus sequences using Racon reduced the number of indels to 0.73% and yielded five indel-free barcodes but this treatment also increased substitution errors in five barcodes. Both consensus sequences were then subjected to the AA-based error correction pipeline.

**Table 4:** Errors rates and number of ambiguities in MinION barcodes as obtained by various methods tested for the three datasets (A, B, C). Mean and range values are provided as % of number of query bps aligned.

| Data-set | Error | MAFFT |  |  | MAFFT+AA |  |  | RACON |  |  | RACON+AA |  |  | CONSOLIDATED<br>BARCODE |  |  |
| --- | --- | --- | --- | --- | --- | --- | --- | --- | --- | --- | --- | --- | --- | --- | --- | --- |
|  |  | N | Mean | Range | N | Mean | Range | N | Mean | Range | N | Mean | Range | N | Mean | Range |
| A | Mismatches | 0 | 0 | 0-0 | 0 | 0 | 0-0 | 5 | 0.03 | 0-0.3 | 5 | 0.005 | 0-1.52 | 1 | <0.001 | 0-0.15 |
|  | Gaps | 471 | 0.76 | 0.15-1.86 | 0 | 0 | 0-0 | 467 | 0.73 | 0-2.2 | 0 | 0 | 0-0 | 0 | 0 | 0-0 |
| B | Mismatches | 0 | 0 | 0-0 | 0 | 0 | 0-0 | 1 | 0.001 | 0-0.25 | 0 | 0 | 0-0 | 0 | 0 | 0-0 |
|  | Gaps | 191 | 1.07 | 0.5-2.05 | 0 | 0 | 0-0 | 181 | 0.92 | 0-1.78 | 0 | 0 | 0-0 | 0 | 0 | 0-0 |
| C | Mismatches | 1 | 0.01 | 0-0.64 | 0 | 0 | 0-0 | NA |  |  |  |  |  |  |  |  |
|  | Gaps | 48 | 2.3 | 1.59-6 | 0 | 0 | 0-0 |  |  |  |  |  |  |  |  |  |

This pipeline corrected the substitution and indel errors for MAFFT consensus barcodes (MAFFT+AA barcodes, Table 4). As expected, the error correction increased the number of ambiguous bases (average of 1.53%; approximately ∼10 bp; Figure 1). For the Racon barcode hypotheses, substitution errors were found in 5 barcodes but all indel errors could be corrected. Overall the percentage of “N” was 1.37% and thus lower than for the MAFFT barcodes. Consolidation of the two barcode hypotheses for each specimen yielded 470 correct barcodes for the 471 with comparison data. All barcodes were free of indel errors, but one retained a substitution error and an average of 1.35% of all bases were ambiguous (ca. 8-9 bp across 658 bp).

**Figure 1:**
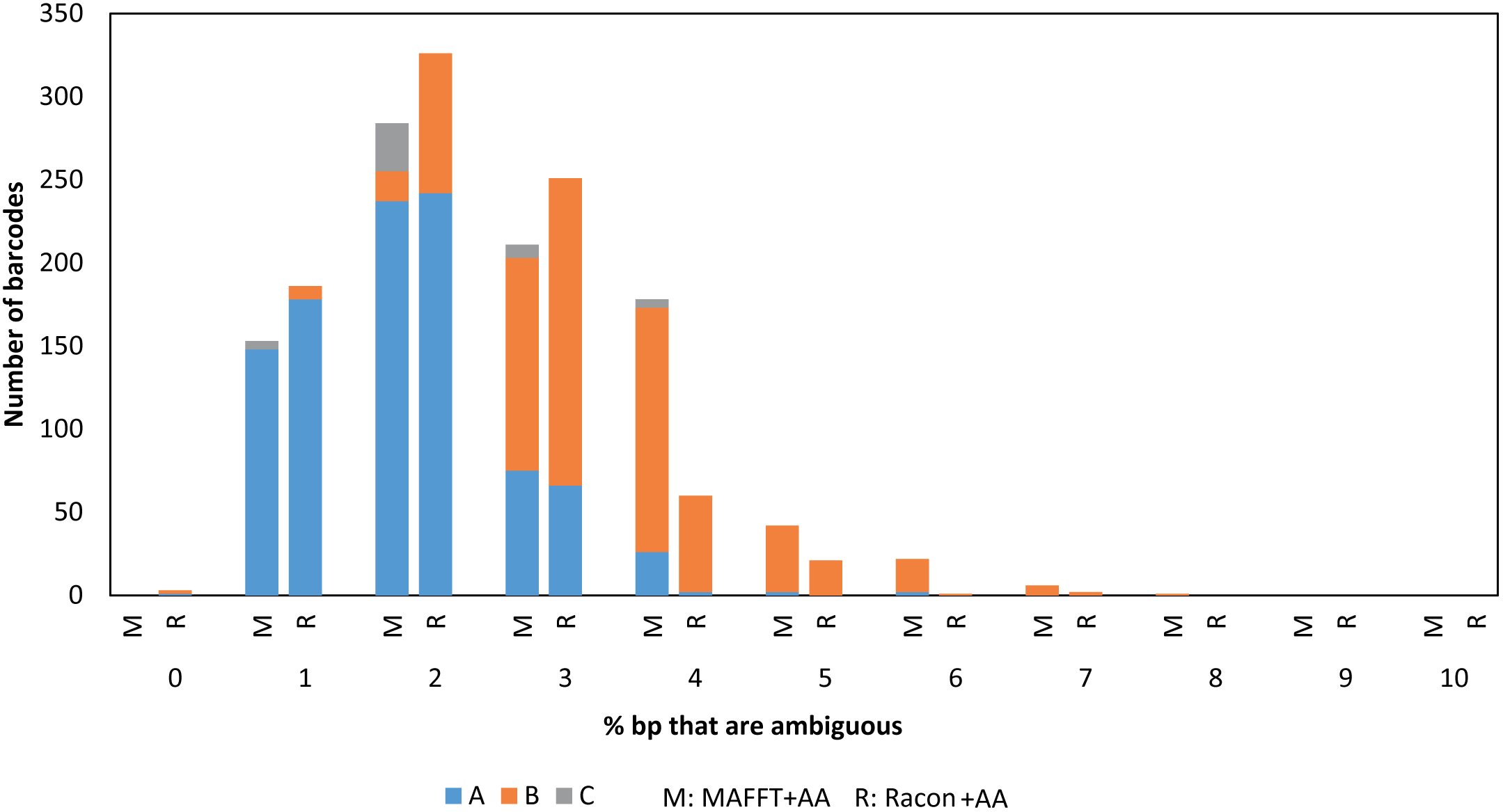
Distribution of % of bases that are ambiguous across different amino acid corrected datasets.

### Validation of the established pipeline

After analysing the main dataset (A), we tested the optimised parameters and methods for the two datasets obtained with different MinION™ chemistries (ants: B; chironomid midges: C). We overall find the performance of the pipeline to be very similar. MAFFT barcodes are substitution-error free but contain indels, MAFFT+AA barcodes contain no substitution and indel errors but are more conservative in that 1.8-3.2% bases are scored as ambiguous. The barcodes obtained by consolidating the corrected MAFFT and Racon barcodes were more accurate than those for Dataset A (no mismatches) and we observe an overall reduction of the number of ambiguous nucleotides.

For datasets B and C, we demultiplexed 428/575 and 50/50 barcodes at 10X coverage from which a clean barcode with <1% ambiguous bases could be obtained for 395 and 48 specimens respectively. In order to validate the results, we obtained 194 and 50 reference barcodes for these two datasets using Illumina MiSeq (see below for details). The comparison of MAFFT and Racon consensus for dataset B showed similar results to dataset A; i.e., substitution errors were marginally higher for Racon consensus (in 0 vs 1/191 barcodes, Table 4) but indel errors lower (1.07% vs 0.92%). Amino acid correction revealed no errors for MAFFT consensus barcodes but the number of ambiguous nucleotides was higher than dataset A at 3.2%. When amino acid correction was applied to Racon consensus barcodes no errors as well as fewer ambiguities were obtained (2.4%). Lastly, the consolidated barcode set contained no substitution/indel errors, and 2.3% ambiguous bases. For dataset C, the one of the 48 MAFFT consensus barcode contained 2 substitution errors. The AA correction pipeline excluded this low coverage barcode (11X) as it contained too many indels while the remaining 47 barcodes were accurate. We did not test consolidation for dataset C as the Racon polishing could not be performed because too few reads were mapped at --max-error 0.05. Note that this dataset was obtained with an outdated chemistry and is here mostly used to test the robustness of our pipelines.

Dataset B had a lower success rate at the demultiplexing stage compared to dataset A, (428/575 DNA barcodes). This reflected the lower PCR success rate for this sample which was estimated based on gel electrophoresis for a subsample to be ca. 74% (∼421 specimens). Using our MinION pipeline, we obtained 395 specimen barcodes using MAFFT with a low proportion of ambiguous bases (N <1%). 359 of these barcodes could be retained after AA correction. This drop of 36 barcodes is due to barcodes that failed the 640-670 length criterion or contained too many indels after alignment with BLAST hits. Several of these 36 aligned poorly, had BLAST hits <80%, but are well supported by raw data with <1% ambiguities and high coverage (>100X). This suggests that a non-functional COI copy was amplified during PCR.

For chironomid midges (C), we sequenced 313 bp barcodes for 50 specimens and obtained 14,772 reads using an R9 MinION™ flowcell. For this dataset we ran our pipeline under the unique index mode and allowed for one mismatch in the tag because we had used shorter 9 bp tags which were not designed for sequencing technologies with high error rates. We obtained >10X coverage for all 50 specimens, while obtaining 48 barcodes (96%) that contained <1% of ambiguous bases

As reference data for these two datasets, we obtained sequences from the same products using Illumina MiSeq. For dataset C, this was a straightforward implementation of the NGS barcoding pipeline of Meier *et al.* (2016), which yielded all 50 barcodes of 313 bp length. The same approach for dataset B yielded clean sequences for both 5’ and 3’ end of the COI barcode for only 194 specimens; i.e., only 400 bp per sequence could be used to assess the accuracy of the MinION barcodes.

### mOTU composition

Overall we obtained four datasets from the three sequencing runs with 243 specimens for Dolichopodidae, 247 specimens for a mixed Diptera sample, 359 specimens for Formicidae, and 47 for Chironomidae. Objective clustering at 3% p-distance threshold revealed identical number of mOTUs from MAFFT+AA and MAFFT DNA barcodes for Dolichopodidae (number of mOTUs=33), mixed Diptera (number of mOTUs=73), Chironomidae (number of mOTUs=3) and Formicidae (number of mOTUs=20). We also assessed whether the reduction of sequence length to 313 bp fragments affected the number of mOTUs. No difference was found for the Dolichopodidae and Formicidae datasets, while the number of species was 74 using the 313 bp fragment for mixed Diptera and 73 for the full length barcodes.

### Effect of run time on sample characterization

MinION™ allows for real-time sequencing and sequences can be analyzed at any point in time. We thus assessed how much data was generated over time and how many specimens are recovered at 10X barcode coverage (Figure 2). For dataset A, 98% of the barcodes could be obtained within 2 hours of sequencing at which point the average coverage was at 100X per barcode (Figure 2 a, b, Supplementary Figure 5). Given the coverage, it is not surprising that the species composition of the sample was stable after 2 hours of sequencing (Bray-Curtis dissimilarity index <0.02 for both Dolichopodidae and mixed Diptera sample, Figure 2c). After 12 hours of sequencing the coverage per specimen reached an average of 467X, following which there were slow improvements to 577X by 48 hours. The slowest accumulation of specimen barcodes was observed for the sequencing run of dataset B (ants); nonetheless after 12 hours, the barcodes for 93% of the specimens could be obtained.

**Figure 2:**
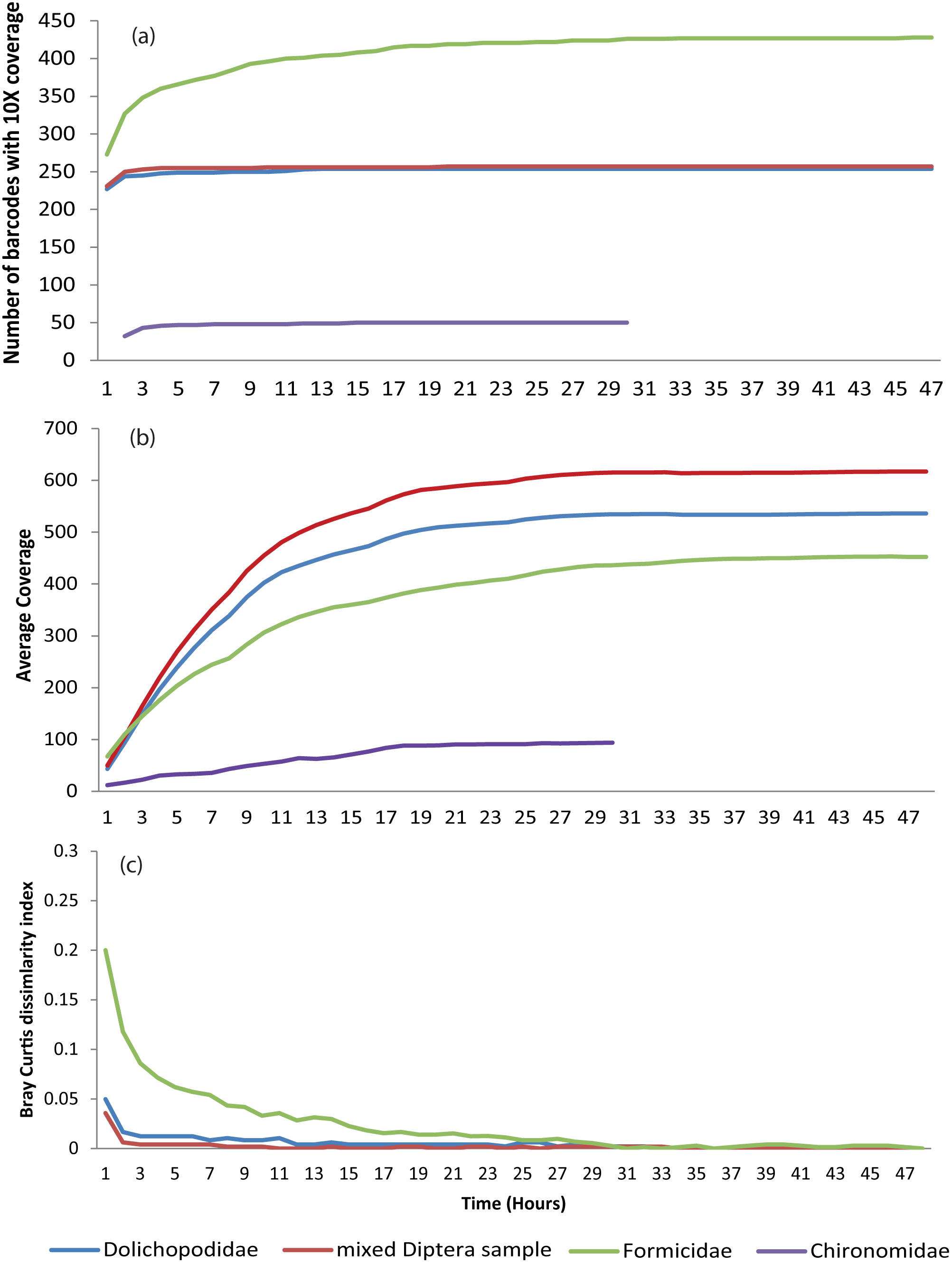
(a) Number of barcodes attaining 10X coverage over time (b) average coverage over time (c) dissimilarity in species composition at a given time point when compared with species composition based on full dataset.

For the largest dataset (A), the average coverage per barcode was >100X within 3 hours of sequencing. At this point only 31% of the data had been generated, but 482 of the 490 barcodes had 10X coverage and passed the threshold of <1% ambiguous bases. Comparison of corrected DNA barcodes at the 3 hour time point with corresponding Sanger barcodes revealed two barcodes with errors (MAFFT+AA). At 6 hours, ∼50% of the data had been demultiplexed, 484 barcodes had obtained a coverage of at least 10X and passed the threshold of <1% ambiguous bases. Overall, three barcodes with errors were found (MAFFT+AA).

## DISCUSSION

We here establish a pipeline for the *de novo* generation of DNA barcodes using Oxford Nanopore’s MinION™ sequencer. We demonstrate that a dual tagging strategy can be used to multiplex >500 specimens using a single flowcell R9.5 MIN107. Our test run yielded 490 DNA barcodes. We were able to compare 471 of all 490 MinION barcodes obtained for this dataset with corresponding Sanger barcodes. This comparison revealed that the MinION barcodes were reliable. We found no substitution errors in the MAFFT barcode hypotheses that can be obtained without reference to external data. The only remaining problems were indel errors which were usually concentrated in homopolymer regions. They were found in all barcodes. Note, however, that the performance of the MinION™ sequencer is here shown to be overall improving because the indel error rate for the latest chemistry/flowcell was lower (0.7%, Racon) than for the two older chemistries that were tested with different datasets. In addition, we demonstrate that our error correction pipeline can neutralize the effect of nearly all of these indel errors by inserting ambiguous bases. Even without using external reference barcodes, an accuracy of 99.3% could be achieved for dataset A, while the use of external data increased the accuracy to >99.9999-100%. This increase in accuracy, however, came at the cost of increasing ambiguous nucleotides to 1.3-1.5%. This raises the question whether such loss of information is likely to affect species identification success rates. Based on our data, this seems unlikely because we show that, even analysing a 313 bp fragment of COI (<50% of full length barcode) is sufficient for yielding species composition estimates that are similar to what is obtained with full-length barcodes. This is indirectly also recognized by the standards established for the “Barcode of Life Data Systems” (BOLD) which requires that a barcode is longer than 500 bp before it can be given a “barcode” status (Ratnasingham & Hebert, 2007). Our MinION barcodes easily meet these targets; >92% of the barcodes have more than 640 bp of information while >99% had >=630 bp of information (consolidated barcodes). On the other hand, a BOLD systems download of the Dolichopodidae COI-5P Sanger barcodes (>=500 bp: 29,914 sequences) revealed that only 24% are >=640 bp long and 29% are >=630 bp long.

We here introduce a simple and effective method for generating MinION barcodes based on commonly-used tools. It consists of aligning reads using MAFFT with subsequent consensus calling. This procedure already yields barcodes that can be used for most identification purposes. For more sensitive applications or for the purpose of submitting barcodes to GenBank, MinION barcodes can be taken through error correction tools such as Racon (Vaser *et al.* 2017), and/or further corrected using the amino-acid correction pipeline proposed and implemented here. Note, however, that even uncorrected MinION barcodes can be used to characterize the species composition of mixed arthropod samples ranging from low complexity samples containing three species to fairly complex samples containing >70 species. We also demonstrate that the data required for generating DNA barcodes can be obtained within 2 hours of sequencing. These are attractive properties for those biologists who need to quickly identify pests, pathogens, vectors, illegally traded species, and verify food ingredients.

### Instrumentation

With regard to instrumentation, MinION™ outcompetes all other sequencing technologies for obtaining barcodes. All barcoding methods share the same instrumentation needs for obtaining amplicons, (pipettes, thermocycler), but the MinION™ sequencer is considerably cheaper and smaller than ABI capillary, Illumina, Ion Torrent, or Roche 454 sequencers. These differences are not trivial because they indirectly also affect how fast barcodes can be obtained: expensive equipment has to be fully utilized in order to be cost-effective and there are usually waiting times for getting access. Moreover handling of expensive instruments requires specialized manpower which in turn increases running cost. In contrast, most laboratories can afford the purchase of multiple MinION™ sequencers (ca. USD 900). The low cost and small size makes the MinION™ also very suitable for establishing temporary laboratories under difficult conditions. Indeed, MinION™ sequencers could be paired with the kind of small thermocyclers that have recently become available and that are suitable for field conditions (Marx, 2015). We estimate that the total equipment cost for a basic field laboratory for barcoding with MinION™ could now be as low as USD 3000 and technologies like the MinION™ will get biologists closer to the vision of being able to obtain sequences in the field. Currently, we note that access to internet and a remote server is required for data analyses due to the computational resources needed for 1D^2^ base-calling (see below).

### Speed

All barcoding procedures involve (a) DNA sequencing and (b) data analysis. In terms of molecular procedures, DNA barcoding requires gene amplification and cleanup of PCR products, irrespective of technology used. For Sanger sequencing this cleanup is specimen-specific while the cleanup procedures for all NGS-based methods (including MinION™) are much faster because they are applied to pooled products. Following clean-up, the MinION™’s library preparation requires <2 hours while Sanger sequencing requires cycle sequencing and cleanup for the individual products. Using liquid handling robots or multichannel pipettes, the latter can be accomplished in 35 minutes for short barcodes (Ivanova *et al.*, 2009), but most protocols require ∼2.5 hours. Capillary sequencing of short Sanger barcodes requires another 45 minutes and is restricted to <=96 samples while the MinION™ sequencer provides species profiles within 2 hours of sequencing and can process 500 amplicons. So, overall the MinION™ and Sanger sequencing require similar amounts of time for small number of samples, but MinION™ is faster than Sanger sequencing for larger numbers of samples. The remaining NGS barcoding protocols (Illumina: Meier *et al.*, 2016; Shokralla *et al.*, 2015; Wang *et al.* 2018; Roche 454: Shokralla *et al.*, 2014) are considerably slower because library preparation and sequencing are more time-consuming.

In terms of data analysis, Sanger barcodes and NGS barcodes differ considerably. Data generated by Sanger sequencing requires chromatogram analyses. There are several tools available that automate this (Sequencher, ASAP (Singh & Bhatia, 2016), etc.). Nonetheless manual inspection of chromatograms is often used to ensure the reliability of base calls and this can be a time consuming effort for 500-1000 barcodes. On the other hand, data analyses of NGS technologies can be automated. With regard to MinION™, the data are generated real time and we were able to make base-calls for a sufficiently large number of reads within ∼2 hours of sequencing based on the latest 1D^2^ chemistry. However base-calling for 1D^2^ reads is computationally intensive and the analyses had to be done on a cluster configured to utilize up to 500 cores. Thus, for field applications it will be necessary to have internet access in order to upload the data onto a server. Fewer computational resources are needed for the next step, i.e., obtaining the “MAFFT barcodes”. For the dataset A (>2 million reads for ca. 500 amplicons), the analysis time using 4 cores was ∼8-9 hours. However, very similar results can be obtained within 1.5 hours on a laptop computer (4 cores, 8 Gb of RAM) by subsampling the read set so that each amplicon is only analyzed based on ∼100 reads. We found that 487/490 barcodes could be called with <1% ambiguity.

### Cost

The cost of sequencing an amplicon with the MinION™ in our latest experiment (∼500 specimens) was approximately USD 1.6/specimen (cost for flowcell: USD 675, reagent costs: USD 130 USD). However, we predict that the number of samples multiplexed in one flowcell could be doubled or tripled because the first 30% of all reads contained the data needed for getting 482 of the 490 barcodes. By the time 50% of the data was generated, 484 barcodes could already be obtained. Note that even at the current cost of USD 1.6, MinION barcodes are substantially cheaper than Sanger barcodes (USD 17/specimen: http://ccdb.ca/pricing/). Nonetheless, the MinION barcodes remain more expensive than NGS barcodes obtained with other NGS technologies (<1 USD: Meier *et al.*, 2016; Hebert *et al.*, 2017). However, the low cost for such barcodes can only be achieved when >10,000 specimens are multiplexed. Overall, we believe that MinION barcodes will soon also be available for <1 USD given that the throughput of MinION™ sequencers has increased >10 fold in one year. Note that the higher throughput was achieved while improving read quality.

### Barcode quality

This remains a drawback of MinION barcodes. The main concern are indel errors that are concentrated in homopolymer sections of the sequences. At a read level, the biases were most prominent in GC-rich homopolymeric regions (Supplementary Figure 6). With our pipeline, many of these errors can be corrected during consensus calling. However given that such homopolymers are prevalent in COI, some errors remain. Fortunately COI is protein-encoding and almost all remaining errors can be detected and resolved when using conserved amino acid motifs for correction. We assessed >390,000 bp of barcode sequence and after correction next to no errors remain; i.e., MinION™ data can probably compete with Sanger data that frequently struggles to detect polymorphisms. However, the low error rate in corrected MinION barcodes come with the cost of having to insert a larger number of ambiguous base calls. For dataset A (ca. 500 barcodes), each full length barcode has ∼8-9 such ambiguous bases (1.3%). We tested whether this leads to a significant erosion in signal, but find no evidence. MinION barcodes can still be unambiguously matched to Sanger barcodes for the same specimen/species. In addition, species composition and abundance estimates based on MinION and Sanger barcodes are identical. There are therefore good reasons to consider the erosion of signal to be a minor concern and we would argue that is better to insert ambiguous bases than to retain indel errors because N’s are less likely to affect the sequence alignment needed for downstream analyses. Note also that the proportion of ambiguous bases is likely to decline. Firstly, we find evidence for an improvement of MinION™ data over time. In addition, the number of ambiguous bases that are inserted during amino acid based correction is dependent on the availability of closely related barcodes in NCBI; i.e., once more barcodes for closely related species become available, fewer ambiguous bases will need to be inserted during the AA correction.

## Conclusions

We here introduce a MinION barcoding pipeline that allows for obtaining DNA barcodes *de novo* without losing the association between sample and barcode. Our basic bioinformatics pipeline is quite straightforward and only requires the input of the FASTA read file obtained from the MinION™, a demultiplexing file specifying the specimen-specific tags, and a few freely available standard alignment tools. Barcodes obtained with these standard tools can be further refined using Racon, AA-correction, and/or consolidation. All these techniques are used to address the main problem of MinION™ reads; i.e., the high error rates in homopolymeric regions. This problem also interferes with demultiplexing because our dual tagging approach required that both forward and reverse tags are identified. This meant that only 14-29% of the reads were retained for analysis for datasets A and B. Using additional tools for increasing this proportion would significantly improve the barcoding capacity of MinION™ flowcells. Overall, we would argue that MinION™ is already suitable for projects of small‐ to moderate scale (<1000 barcodes). For these, the method is attractive because it allows for the rapid turnaround of time-sensitive samples for which the barcode-to-sample association has to be maintained. This association is important for food authentication (Shokralla *et al.*, 2015a), but similarly desirable for bioassessment and the identification of invasive species. For these purposes, it is important that a pipeline allows for the identification of unexpected pests and invasives for which the publically accessible barcode databases lack reference sequences.

## ACKNOWLEDGEMENTS

The project was supported by funding from the South East Asian Biodiversity Genomics Center (NUS grant nos. R-154-000-648-646 and R-154-000-648-733) and from A*STAR. We thank Dr. Sujatha N. Kutty and Arina Adom for testing scripts.

## AUTHOR CONTRIBUTIONS

RM, NN and AS designed the experiments, BB, WW, WT, AHQN and EJHB did the molecular work, AS developed the pipeline with input from RM, AS, KJY and DB analysed the data. AS and RM wrote the manuscript with inputs from all authors.

